# Limitations of Busulfan to Create Humanize Mice with an Innate Immune System

**DOI:** 10.1101/2022.07.15.500220

**Authors:** Ohud Alhawiti, Bin Hu, Jennifer Kobilinski, Chunqing Guo, Joseph W. Landry

**Affiliations:** Department of Human and Molecular Genetics, Institute of Molecular Medicine, Massey Cancer Center, Virginia Commonwealth University School of Medicine, Richmond, VA, 23298; Department of Pathology, Mouse Models Core, Virginia Commonwealth University School of Medicine, Richmond, VA, 23298

## Abstract

Humanized mouse models have improved biomedical research by providing a tractable system with which to perform *in vivo* experiments on human tissues. Use of irritators is the standard method for establishing high levels of stem cell engraftment, however not all institutes have access to this instrumentation in the animal facility. The use of busulfan has been successfully used to precondition for stem cell engraftment on a limited number of mouse backgrounds. In this report we further test the utility of busulfan as a means to successfully engraft hIL15-Tg-NSG and SGM3-NSG mouse stains which are capable of establishing the innate NK cell and myeloid immune compartments. Results from our studies show that busulfan can successfully precondition hIL15-Tg-NSG mice but not SGM3-NSG mice for high levels of human immune cell engraftment. SGM3-NSG mice preconditioned with busulfan exhibited only 10-20% human CD45 cells in the bone marrow or spleen, where as NSG and hIL15-Tg-NSG mice routinely achieved ∼80%. Busulfan preconditioned SGM3-NSG mice showed elevated levels of granulocytic MDSC, and cDC1 and cDC2 myeloid populations. This is in contrast with hIL15-TG-NSG which showed robust reconstitution of mature CD16 expressing NK cells. We conclude from our studies that busulfan is an effective means to precondition mice for CD34+ stem cell engraftment, but it may have limitations when use to precondition the SGM3-NSG model.

## INTRODUCTION

Increasingly biomedical research is utilizing humanized mouse models to study human cells, tissues, and organs in a xerograph setting (Stripecke et al. 2020). In theory, these models allow the study of human immunology in mice without the ethical hurdles involved with experimentation on human patients. Since the first mice with a humanized immune system were developed over 20 years ago (Belizario 2009), there has been a rapid expansion of available mouse strains, techniques, and methods of analysis to use these mouse models (Yip et al. 2019). More so than other disciplines immunology has benefited greatly from humanized mouse models (Ishikawa et al. 2005a; Ito et al. 2012). In this capacity, mice with a humanized immune system are used to study a variety of immune diseases including drug toxicity to immune cells, graft vs host disease (GVHD), allergies, immune response to pathogens, the anti-tumor immune response, and in the development of immunotherapies (Hyochol Ahn, PhD, Michael Weaver, PhD, Debra Lyon, PhD, Eunyoung Choi, RN, and Roger B. Fillingim and Tumbar 2017; Shultz, Ishikawa, and Greiner 2007). Most recently, significant efforts are being made to develop more humanized models to test and study checkpoint inhibitors (Yip et al. 2019).

Humanization of a mouse immune system can occur through the transplantation of immune organs (fetal liver, thymus, etc…), peripheral blood cells (PBC), or through the engraftment of CD34^+^ hematopoietic stem cells (Martinov et al., 2021). Commonly, the humanized immune system is made using NOG (NOD/Shi-scid, Il2rg−/−), NSG (NOD/LtSz-scid, Il2rg−/−), or the BRG (Balb/c Rag2−/−, Il2rg−/−) backgrounds (Ishikawa et al. 2005a; Ito et al. 2012). These models lack mature murine T, B and natural killer (NK) cells and have compromised macrophages, and when engrafted with human immune cells are superior for promoting the formation of human hematopoietic cells via physiological developmental intermediates (Audigé et al. 2017). When reconstituted with human CD34^+^ stem cells these mice will develop the most basic components of the human immune system including B cells, CD8 and CD4 T cells, and undifferentiated monocytes (Ishikawa et al. 2005b). Largely due to deficiencies cytokines, humanized NSG, or equivalent background, lack key immune cells including NK cells, T regulatory cells (Treg) and mature differentiated myeloid linage, to name a few, (Audigé et al. 2017). As such, these humanized mice can mimic some human immunological responses, such as antibody formation in response to antigens and T cell cytotoxicity (Shultz, Ishikawa, and Greiner 2007), but largely lack functional innate immune cells. The limited reconstitution of some cells of the immune system in mice has incentivized the development of additional mouse models expressing the missing, but critical, immune promoting cytokines. NSG mice expressing human IL15 (hIL15Tg-NSG) can support human NK cell growth, maturation, and function. As such it is be used to examine human NK cell biology and NK-mediated cancer immunotherapy (Ju et al. 2019). NSG mice expressing transgenic human SCF, GM-CSF and IL3 (SGM3-NSG) mice promote the growth of mature differentiated myeloid and Treg cells (Middlebrook et al. 2017). When humanized both the hIL15Tg-NSG and SGM3-NSG mice can be a crucial tool for studying human immune cell biology and assessing the efficiency of new therapies in the lab before they are brought to the clinic (Norman P Spack Daniel E Shumer 2017).

Use of humanized mouse models require significant investment in infrastructure required for engraftment (irritator) and the long-term breeding and housing of severely immune compromised mouse models (ultra-barriers) (Belizario 2009). Total body irradiation (TBI) has been a standard conditioning regimen to achieve high levels of human cell engraftment in these mouse models (Belizario 2009). However, the TBI procedure requires the use of expensive and dedicated irradiators, which must be located inside the barrier facility, and therefore limit many institutes without access to this infrastructure. Because of this, researchers have found a way to replace TBI with chemotherapeutic drugs like the bone marrow ablative chemotherapy busulfan (Hayakawa et al. 2009).

In order to establish simple humanization process with the ability to reconstitute T cells, B cells, and the more challenging NK and mature differentiated myeloid immune cell compartments, we developed a simple method using busulfan conditioning followed by transplant of CD34+ cells into the NSG, hIL15Tg-NSG and the SGM3-NSG mouse backgrounds.

## MATERIALS AND METHODS

### Animals

Breeder pairs of (NSG) NOD.Cg-Prkdc^scid^ Il2rg^tm1Wjl/SzJ^ (Strain #005557), (SGM3-NSG) NOD.Cg-Prkdc^scid^ Il2rgtm1Wjl Tg(CMV-IL3,CSF2,KITLG)1Eav/MloySzJ (Strain #013062) and (hIL15Tg-NSG) NOD.Cg-Prkdc^scid^ Il2rg^tm1Wjl^ Tg(IL15)1Sz/SzJ (Strain #030890) at six weeks age were purchased from the Jackson Laboratory. The mice were bred, housed and all procedures were performed in the Massy Cancer Center ultra-barrier facility at Virginia Commonwealth University (Richmond, USA) under specific free pathogen conditions. Trials were performed under our institution’s guidelines for the ethical use of animals under IACUC protocol AD10000017 and its modifications.

### Human CD34^+^ Cell Purification

Cryopreserved and deidentified human umbilical cord blood samples (hUCB) were obtained from Duke University Stem Cell Repository under a VCU IRB exempt protocol. Briefly, 22 ml hUCB was diluted 1:1 into 10% BSA PBS until a final dilution of 1:8 was achieved. Cells were centrifuged at 500 x g for 10 min and suspended in 10 ml red blood cell (RBC) lysis buffer. After RBC lysis the cells were resuspended in 8 ml of 10% BSA PBS and layered over 8 ml of Ficoll Plaque Plus Premium then centrifuged at 400g for 30 minutes. Mononuclear cells (MNCs) were removed from the gradient interface, washed, and then suspended in 10% BSA PBS. Cells were counted and purified using a human CD34+ stem cell purification kit (Miltiney Biotech catalog #130-100-453) according to manufactures protocol. Human CD34^+^ stem cell purity was monitored by anti-human CD34 FITC (clone:581) and 7-AAD viability dye using LSR Fortessa X-20 flow cytometer (BD Biosciences).

### Mouse Conditioning and Engraftment

Busulfan (Fisher Scientific, Cat# 11-101-7872) was dissolved in DMSO and diluted with 0.9 percent saline to obtain of 5 mg/ml busulfan in 30% DMSO. The busulfan solution was administered IP at 25 mg/kg to mice 48 and 24 hours before transplantation. The day after the last busulfan treatment, human CD34^+^ cells were transplanted via IV injection in 100 µL PBS. The number of stem cells used varied depending on the experiment and are noted in the legend of each experiment. Mice were monitored and weighed weekly to monitor for GVHD.

### Engraftment Analysis

Blood samples were drawn from the tail at designated times, treated with K_2_EDTA (2 µl 75% K_2_EDTA /100 µl blood), processed by using RBC lysis buffer, and finally washed in PBS. Standard techniques were used to prepare cell suspensions from the spleen and bone marrow. Cells were stained with Zombie Aqua for viability, Fc blocked, then stained with combinations of the following antibodies: BV711 anti-human CD8α (clone:RPA-T8), APC-Fire750 anti-human CD11c (clone:3.9), PE anti-human CD19 (clone:HIB19), AF488 anti-human CD3 (clone:OKT3), PE-Cyanine5 anti-human CD4 (clone:OKT4), AF700 anti-human CD4 (clone:OKT4), FITC antihuman CD34 (clone:581), APC anti-human CD56 (clone:5.1H11), PE/Cyanine7 anti-human CD45 (clone:2D1), BV421 anti-human CD33 (clone:WM53), FITC anti-human CD16 (clone:3G8), PE anti-human CD40 (clone:5C3), BV421 anti-human perforin (clone:G9), PE anti-human CD69 (clone:FN50), PE-Cyanine5 anti-human CD14 (clone:61D3), APC anti-human CD141 (clone:M80), FITC anti-human CD204 (clone:9E4A8), BV605 anti-human HLA-DR (clone:L243), and BV711 anti-humanCD1c (clone:L161). Stained samples were post fixed and analyzed by the BD LSR Fortessa X-20 flow cytometer.

## RESULTS

### Engraftment of NSG Mice with hUCB-Derived CD34^+^ Cells using Busulfan

In **Figure 1A** we outline our methodology for creating humanized mice at Virginia Commonwealth University. NSG mice were treated IP with 25 mg/kg body weight busulfan at 48 and 24 hours before receiving purified hUCB CD34^+^ stem cells by IV. Our CD34^+^ stem cell purification protocol, which uses a Ficoll density gradient centrifugation, followed by positive selection purification of CD34^+^ cells typically achieved 96.7%-98% (N=3) purity in our human CD34^+^ cell preparations from hUCB samples **(Fig. 1B)**. An assessment of the effectiveness of preconditioning of the NSG mice for CD34+ engraftment shows that busulfan treatment (at day -2 and -1) is required to achieve engraftment of the hCD45+ cell population **(Fig. 1C)**.

**Figure 1.).**
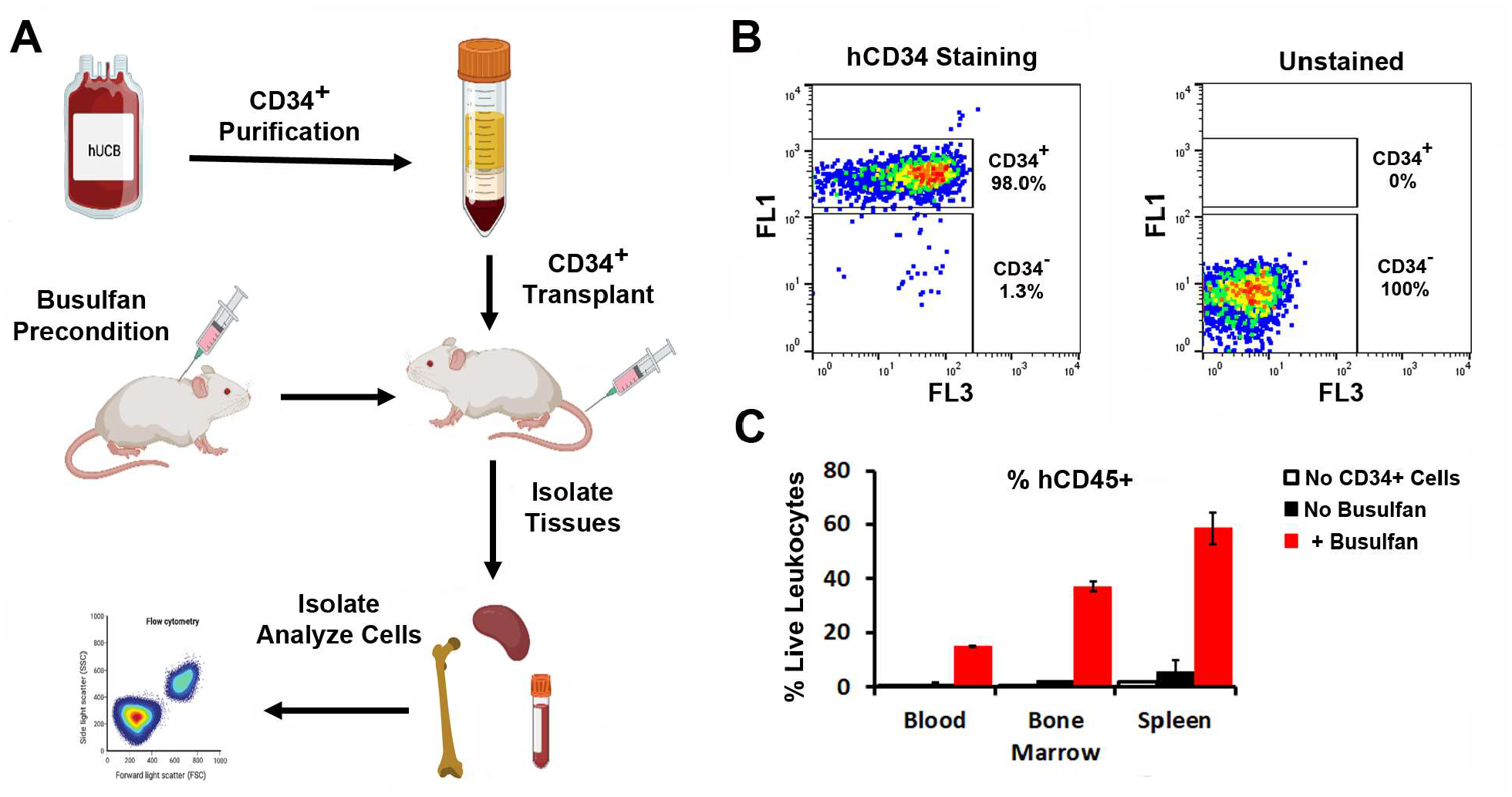
Scheme for Constructing Humanized Mice. **(A)** CD34^+^ cells were isolated from hUCB using a Ficoll-Paque gradient followed by an Ultrapure Human CD34 MicroBead Kit from Miltiney Biotech. Busulfan was used to condition the mice 48 and 24 hrs before transplanting purified CD34^+^ cells via IV injection. Mice were housed in an ultra-barrier facility until experimental endpoint was achieved. The humanized blood, bone marrow, and spleen were removed and analyzed using a flow cytometer at set time points. (**B)** Analysis of representative human CD34^+^ cell purification from hUCB using anti-human CD34^+^ and 7AAD by flow cytometry. **(C)** Effectiveness of busulfan treatment for the engraftment of human CD34+ cells and expansion of hCD45+ cells from in the blood, bone marrow and spleen was determined in NSG mice. N=2 per group.

In our first experiment we transplanted 10^5^ CD34^+^ cells into busulfan conditioned NSG mice. Mice were monitored over the course of the experiment and no signs of GVHD (includes >15% weight loss, hunched posture, fur loss, reduced mobility and tachypnea) over 12 weeks were observed **(Fig. 2A)**. At the 15^th^ week post-transplant, human immune cell engraftment was monitored from blood, spleen, and bone marrow samples by flow cytometry (**Fig. S1**). As a control we performed a similar analysis on a control NSG mouse which was treated with busulfan but did not receive human CD34^+^ cell transplant. From this control we did not detect any human CD45^+^ (hCD45^+^) cells, suggesting that our staining was specific for human cells (**Fig. 2B**). Our assessment of human immune cells observed ∼25%, ∼78% and ∼85% hCD45^+^ cells from total live leukocytes in the blood, bone marrow and spleen, respectively, for busulfan conditioned mice transplanted with CD34^+^ cells. Our ability to reconstitute ∼25% hCD45^+^ immune cells in the blood is approximate to what was previously reported for using busulfan (Kang et al. 2016).

**Figure 2.).**
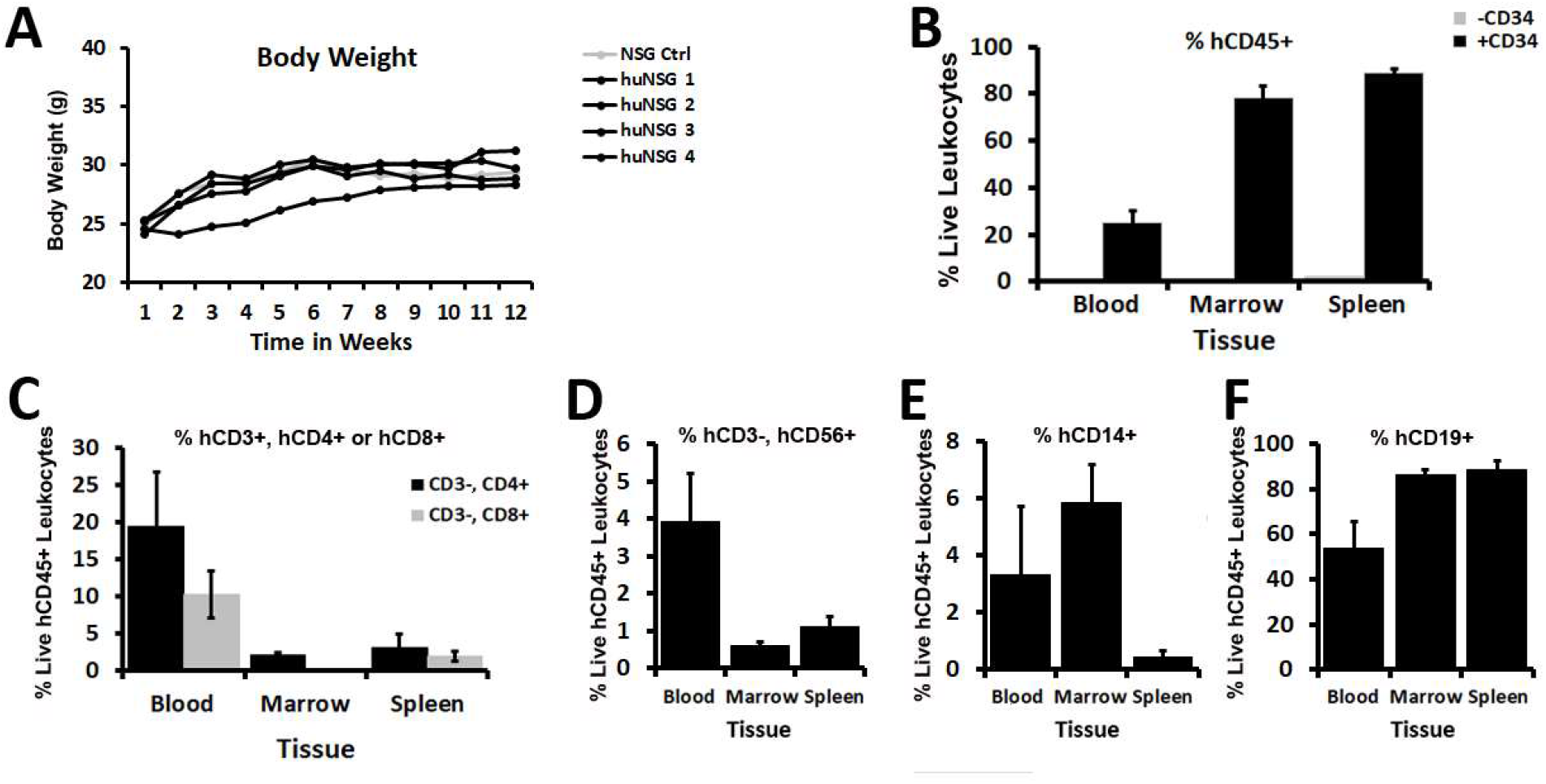
Using Busulfan to Humanize the Immune System of NSG Mice. **(A)** The body weight of humanized NSG mice were measured once a week after engraftment. (**B-F)** The blood, spleen and bone marrow of NSG mice 15 weeks post transplantation with hUCB-derived hCD34^+^ cells used were analyzed for abundance of (**B**) hCD45+ human leukocytes, (**C**) hCD3+, hCD4+ or, hCD3+, hCD8+ human T cells, (**D**) hCD3-, hCD56+ human NK cells, (**E**) hCD14+ human monocytes or (**F**) hCD19+ human B cells.

In addition to bulk hCD45^+^ immune cells we monitored specific human immune cell subsets included CD4 T cells (hCD45^+^, hCD3^+^, hCD4^+^), CD8 T cells (hCD45^+^, hCD3^+^, hCD8^+^), NK cells (hCD45^+^, hCD3^-^, hCD56^+^), monocytes (hCD45^+^, hCD14^+^), and B cells (hCD45^+^, hCD19^+^) from the blood, bone marrow and spleen (**Fig. 2C-F**). As expected, we observed an enrichment of CD4 and CD8 T cells (3:1 ratio CD4^+^ to CD8^+^) and a low number (∼4% to 0.5% of live hCD45^+^) NK cells in the blood, bone marrow and spleen. CD14^+^ monocytes were observed at ∼6% to 0.5% of live hCD45^+^ cells primarily in the blood and the bone marrow. As previously reported (Choi et al. 2011), we observed that the majority of human immune cells in the reconstituted mice were B cells (hCD45^+^, hCD19^+^) which were present in abundance at ∼50%, ∼90% and ∼90% of live hCD45^+^ cells in the blood, bone marrow and spleen (**Fig. 2F**). This pilot study demonstrated that the use of busulfan can result in the engraftment of human immune cells, effectively leading the humanization of NSG mice.

### The Use of Busulfan to Humanize hIL15Tg-NSG and SGM3-NSG Mice with hUCB-Derived CD34^+^ Cells

Previous reports have shown that busulfan can successfully humanize NSG mice with CD34^+^ stems cells (Kang et al. 2016), which we have confirmed in our studies (**Fig. 2**). To expand on these findings, we asked if busulfan can precondition specialty strains of NSG mice for the enrichment of specific human innate immune cells populations.

To create mouse models with humanized NK cells, we busulfan treated hIL15Tg-NSG and SGM3-NSG mice using the same protocol as described (**Fig. 1A**). Three controls were performed in this experiment. The first, we reconstituted NSG mice (hIL15Tg-NSG or SGM3-NSG) to determine the impact of the hIL15Tg-NSG background on NK cell reconstitution, and the SGM3 background to study myeloid lineage reconstitution. The second control we included donor sex as a biological variable in our design. In our design we used a single male donor to reconstitute 3 male NSG and 3 hIL15TG NSG mice, and a single female donor to reconstitute 3 female NSG and 3 SGM3 mice. We did not include into our design transplanting male or female hUCB into mice of the opposite sex, because of incompatibility of the sex hormones (Taneja 2018). With this design we can determine the impact of the donors sex on the reconstitution of the NSG mice. As a final control we included a busulfan treated NSG mouse which was not transplanted with CD34^+^ stem cells as a control for our antibody staining.

We first compared the ability of the male and female donors to reconstitute the innate lineages including both mature and activated NK cells, and the development of mature and activated AP myeloid cells. From this analysis, we observe equivalent human immune cell reconstitution with ∼10%, ∼60%, and ∼60% hCD45^+^ cells from total live leukocytes in the blood, bone marrow and spleen, respectively, between the 2 donors in the NSG background (**Fig. S2**). Note that one NSG mouse reconstituted with the male donor did not have any hCD45^+^ cells suggesting a failed reconstitution, and it was removed from the analysis (data not shown). Further analysis shows that the monocyte population (hCD45^+^, hCD14^+^) at ∼7-3% for both sexes are equivalent in the blood, bone marrow and spleen. A similar equivalent was observed for the NK cell population (hCD45^+^, hCD56^+^) which showed ∼2%-0.5% for both donors in the blood, bone marrow and spleen (**Fig. S2**). From these results we conclude that both male and female donors can reconstitute busulfan conditioned mice equivalently and have pooled their data for the remainder of the analysis.

As with the studies on the NSG background from pilot studies in **Figure 2** both the NSG controls and the hIL15Tg-NSG mice were monitored over the course of the experiment and no signs of GVHD (includes >15% weight loss, hunched posture, fur loss, reduced mobility and tachypnea) over 12 weeks were observed (**Fig. 3A**). In the first comparison, we examined the hCD45^+^ population in NSG and hIL15Tg-NSG mice in blood, bone marrow, and spleen 15 weeks after engraftment. Note that because we did not observe any differences in the sex of the donor in the reconstitutions of human immune cell populations (**Fig. S2**), we aggregated both donor data from the NSG reconstitution (N=5), and compared to hIL15Tg-NSG (N=3). From this analysis, we show that the average percent hCD45^+^ (hCD45^+^ of total leukocytes) cells was greater in the blood in hIL15Tg-NSG mice compared to NSG mice (∼23% vs 10%)(**Fig. 3B**). In the bone marrow and the spleen, we observed equivalent reconstitution with ∼70% hCD45^+^ in both hIL15Tg-NSG mice compared to NSG (**Fig. 3B**).

**Figure 3.).**
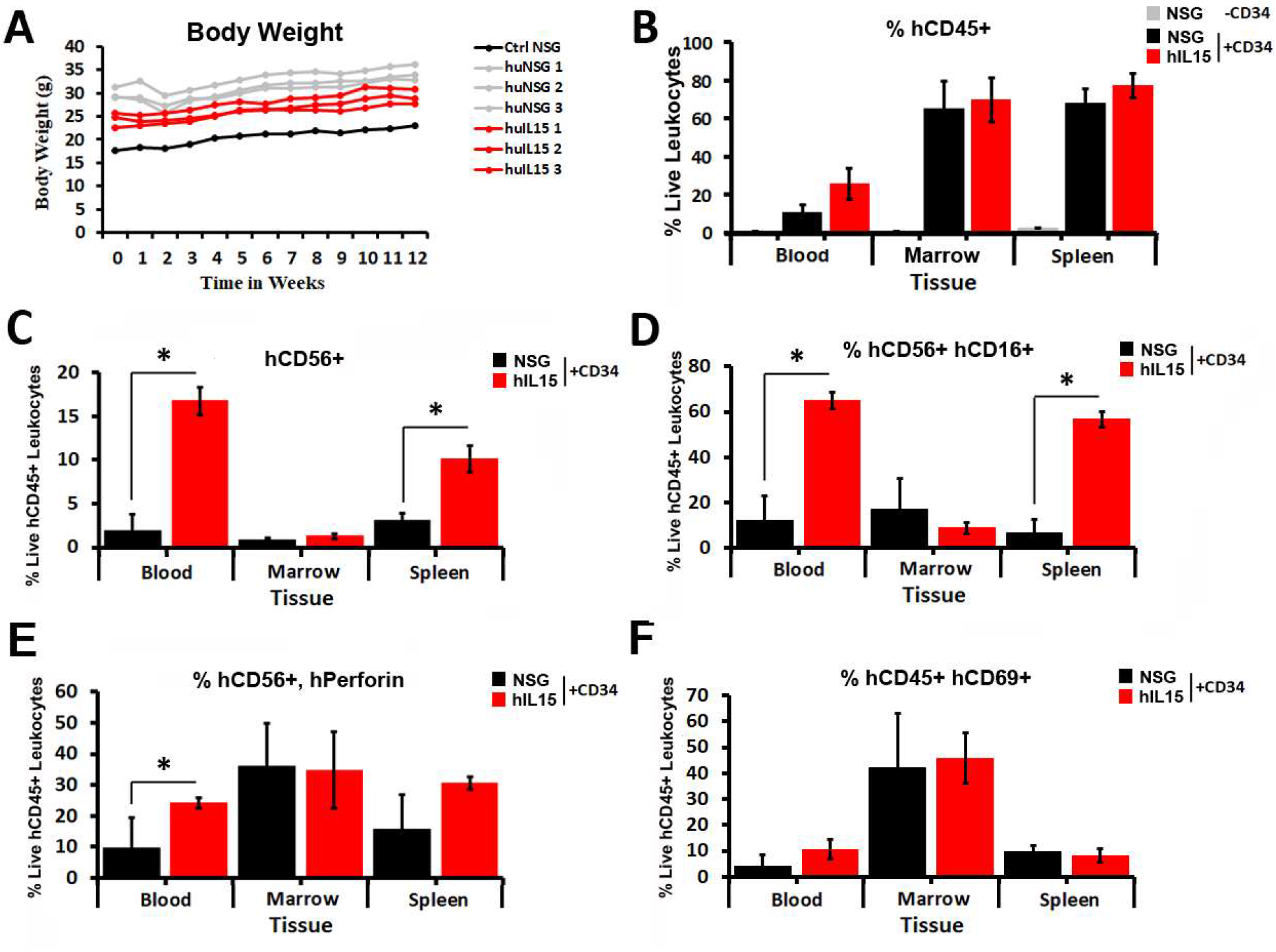
Using Busulfan to Humanize the Immune System of hIL15-Tg-NSG Mice. **(A)** The body weight of humanized IL15Tg-NSG and NSG mice were measured once a week for 12 weeks after engraftment. (**B-F)** The blood, spleen and bone marrow of IL15Tg-NSG and NSG mice 15 weeks post transplantation was analyzed for abundance of (**B**) hCD45^+^ human leukocytes, (**C**) hCD56^+^ human NK cells, (**D**) hCD56^+^ hCD16^+^ mature human NK cells, (**E**) hCD56^+^ perforin^+^ mature human NK cells, (**F**) hCD56^+^, hCD69^+^ active human NK cells.

To assess the ability of the hIL15Tg-NSG mice to reconstitute human NK cells we next measured NK cell abundance, maturation and activation in blood, bone marrow and spleen. The NK cell abundance (hCD45^+^, hCD56^+^) was higher in the blood and spleen in hIL15Tg-NSG mice compared to NSG mice, ∼15% to ∼3%, and ∼10% to ∼3%, respectively (**Fig. 3C**). NK cell maturation was also higher in hIL15Tg-NSG mice compared to NSG mice in these same locations. We observed ∼60% of the human NK cells to express the NK cell maturation marker hCD16 in the blood and spleen, whereas only ∼10% express the same marker in these same locations in NSG mice (**Fig. 3D**). An analysis of the NK cell maturation marker perforin shows a similar trend with elevated perforin expressing NK cells in hIL15Tg-NSG mice in both the blood and spleen, but not to the same degree of significance (**Fig. 3E**). Staining of the hCD69 activation marker was equivalent in the blood, marrow and spleen between the hIL15Tg-NSG and NSG mice (**Fig. 3F)**. These results in total show that busulfan can be used to create humanized IL15Tg-NSG mice which reconstitute a significant improvement in mature human NK cell compartment.

Continuing our assessment of busulfan’s ability to reconstitute the human immune system in specialty NSG strains mice we analyzed the reconstituted SGM3-NSG mice. As observed with the NSG (**Fig. 2**) and hIL15Tg-NSG mice (**Fig. 3**) we did not observe any evidence of GVHD over the course of the experiment (**Fig. 4A**). In contrast to hIL15Tg-NSG mice, which reconstitute improved (blood) or roughly equivalent levels of human hCD45^+^ immune cells (bone marrow and spleen), SGM3-NSG mice have significantly lower human hCD45^+^ immune cells than NSG mice in blood, bone marrow and spleen (**Fig. 4B**). These results are not in agreement with published reports which showed enhanced human immune cell engraftment in SGM3-NSG mice when conditioned with radiation (Martinov et al. 2021). We next assessed the maturation and activation status of myeloid cells in the humanized SGM3-NSG mice. We first analyzed for the reconstitution of myeloid derived suppressor cells (MDSC), an important immune suppressive immature monocyte lineage (Audigé et al. 2017). From this analysis we observed increased MDSC cells in the blood, but no significant enrichment of this cell type in either the spleen or bone marrow in either strain, of SGM3-NSG mice compared to NSG mice (∼40% vs ∼8%) (**Fig. 4C**). Additional analysis of hCD14 on the MDSC population segregated them into the monocytic, compared to the granulocytic, MDSC lineage (**Fig. 4D**). From this analysis we observed elevated granulocytic MDSC (hCD14^-^) in the blood, spleen and bone marrow of SGM3-NSG mice compared to NSG mice. As a measure of mature antigen presenting cells (APC), we monitored the reconstitution of cells expressing the dendritic cell markers hCD11C and HLA-DR (**Fig. 4E**). From this analysis we observed decreased levels of cells with dendritic cell markers in the blood in SGM3-NSG mice compared to NSG mice (∼0.1% vs 1.5%), but increased abundances of cells with these same markers in SGM3-NSG mice in the bone marrow and spleen (**Fig. 4E**). To determine the reconstitution of dendritic cell subtypes, and their activity, we monitored for the expression of differentiation markers hCD1c, hCD141, hCD209 and activation marker hCD40 on the CD11c^+^, HLA-DR^+^ population (**Fig. 4F-I**). From these analyses we observe consistently higher abundance of conventional dendritic cell 1 (cDC1) marker hCD141^+^ and cDC2 marker hCD1c^+^ on dendritic cells in the spleen of SGM3-NSG mice compared to NSG mice, but no difference in the abundance of monocytic dendritic cell marker (mo-DC) hCD209^+^ cells in any tissue (**Fig. 4F,H**). The observed abundance of the dendritic cell populations is approximate to that observed in human tissues with the cDC1 and cDC2 being ∼1% or less of the live hCD45^+^ population (Refs). The activation of the dendritic cell population as measured by hCD40 expression was observed to be higher in the bone marrow and spleen of SGM3-NSG mice compared to NSG mice (**Fig. 4I**). The enhanced activation of the DC populations suggests that they may be competent to present antigens to activate human T cells.

**Figure 4.).**
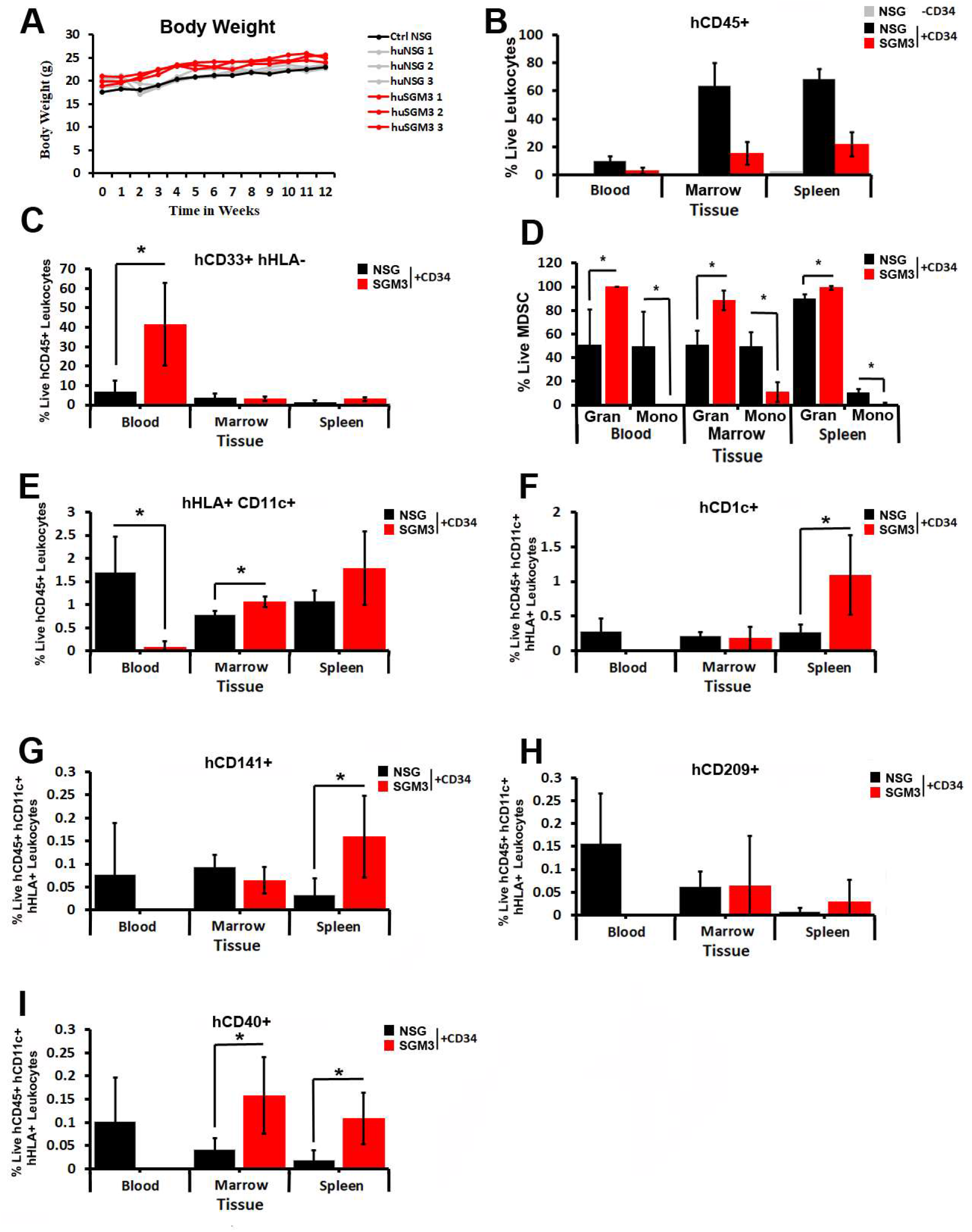
Using Busulfan to Humanize the Immune System of SGM3-NSG Mice. (**A**) The body weight of humanized SGM3-NSG and NSG mice were measured once a week for 12 weeks after engraftment. (**B-I**) The blood, spleen and bone marrow of SGM3-NSG and NSG mice 15 weeks post transplantation was analyzed for abundance of (**B**) hCD45^+^ human leukocytes, (**C**) hCD33^+^ hHLA-DR^-^ human MDSC, (**D**) separation of MDSC into granulocytic (hCD14^-^) and monocytic (hCD14^+^) populations, (**E**) hCD11c^+^ hHLA-DR^+^ dendritic cells, (**F-I**) measurement of hCD1c+, hCD141+, hCD209+ DC populations and (**I**) hCD40+ active dendritic cell populations.

### Determination of Minimal Number CD34+ Stem Cells for Effective Reconstitution

In order for humanized mice to be used in large scale experiments it is essential that they can be reconstituted with a minimum number of purified CD34^+^ stem cells. To this end we wanted to determine the relationship between the number of CD34^+^ cells transplanted and the number of CD45^+^ immune cells observed in the blood over time and if a reduced number of stem cells can be transplanted which can eventually reconstitute the mouse to high proportion hCD45^+^ immune cells. For this experiment we busulfan treated hIL15-Tg-NSG mice which were then transplanted 1.0 × 10^5^, 3.0 × 10^4^ and 1.0 × 10^4^ CD34^+^ stem cells using our standard method **(Fig. 1A)**. We observed slight weight loss in one of the mice reconstituted with 3.0 × 10^4^ from weeks 10 to 14, which rebounded back after week 16 (**Fig. 5A**). The reason for the weight loss is unknown as it was not accompanied with any other symptoms of GVHD (data not shown). Human immune cell abundance was measured from blood draws at 8, 10, 14 and 16 weeks. The average human CD45^+^ cells at week 8 post transplantation were ∼10%, ∼6% and ∼2% for 1.0 × 10^5^, 3.0 × 10^4^ and 1.0 × 10^4^ CD34^+^ stem cells transplanted (**Fig. 5B**). On average there was a minimal increase in hCD45^+^ immune cells over the time interval suggesting that higher numbers of stem cells are needed for effective reconstitution to 16 weeks. The abundance of CD4 (hCD45^+^, hCD3^+^, hCD4^+^), CD8 (hCD45^+^, hCD3^+^, hCD8^+^) T cells, NK cells (hCD45^+^, hCD3^-^, hCD56^+^), and monocytes (hCD45^+^, hCD14^+^) remained consistent as a percentage of CD45^+^ cells over weeks 8-14 (**Fig. 5C-G**). However, at week 16 we observed a significant decrease in B cells and CD4 T cells and an increase in CD8 T cells and NK cells (**Fig. 5C-G**).

**Figure 5.).**
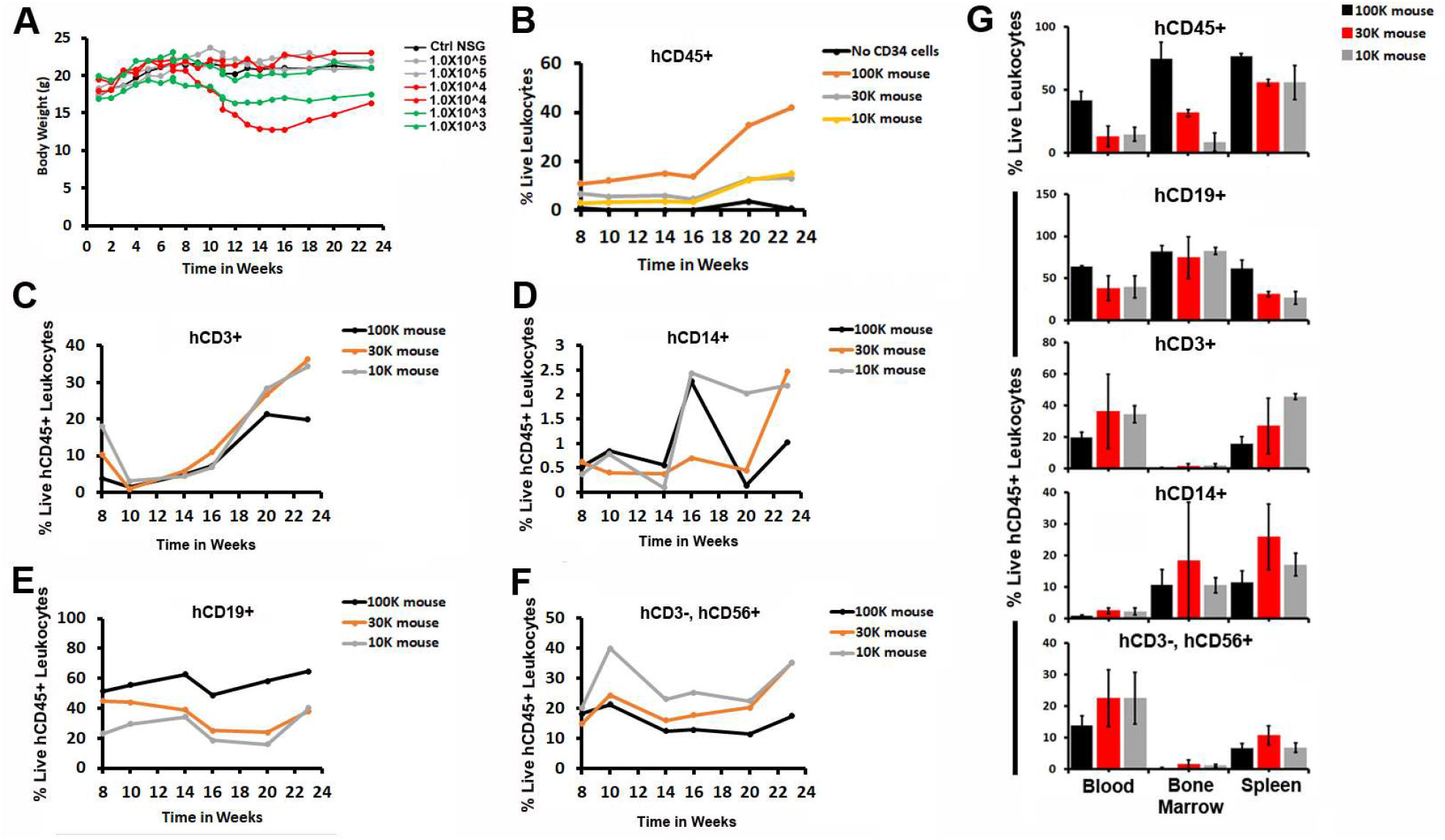
Minimum Number of CD34+ Stem Cells Required for Effective Reconstitution. **(A)** The body weight of humanized IL15Tg-NSG mice reconstituted with 100k, 30k, or 10k CD34^+^ cells were measured from the blood over a 23 week period after engraftment. (**B-G**) The blood of humanized IL15Tg-NSG mice were analyzed for abundance of (**B**) hCD45^+^ human leukocytes, (**C**) hCD3^+^ T cells, (**D**) hCD14^+^ human monocytes, (**E**) hCD19^+^ human B cells (**F**) hCD3^-^, hCD56^+^ human NK cells. **(G)** Analysis of hCD45+, hCD19+, hCD3+, hCD14+ and hCD3-, hCD56+ populations from the blood, bone marrow and spleen at 23 weeks.

## DISCUSSION

Humanized mice are increasingly being used in biomedical research (Kang et al. 2016). These models allow the study of *in vivo* human biology taking advantage of the relative ease and convenience of the mouse model. These models have many uses including the study of drug action on human immune cells, the study of human immune diseases and the anti-tumor immune response to human tumor models (Drake, Chen, and Chen 2012). Specialty strains of mice are being developed rapidly to improve humanization, and immune cells function once established. Examples include the hIL15Tg-NSG and the SGM3-NSG strains, which have been reported to improve NK cell and myeloid cell development (Martinov et al. 2021). Preconditioning of these specialty strains for CD34^+^ engraftment has traditionally used TBI (Kang et al. 2016) to yield high levels of stem cell engraftment (Quesenberry et al. 1999). However, this requires costly irradiators dedicated to barrier animal facilities, which many research institutes do not have available. As such, we set-out to test if the bone marrow depleting drug busulfan could replace TBI as a method to engraft hIL15Tg-NSG and the SGM3-NSG strains for the reconstitution and study of the human innate immune system.

Our initial studies using busulfan to reconstitute the commonly utilized NSG background yielded results similar to those reported in the literature. Over several experiments humanizing NSG mice with CD34^+^ cells we observed no significant change in body weight and no signs of GVHD, which is consistent with the reported success rates in the literature (Kang et al. 2016). Typically, we observe between 10-25% hCD45^+^ cells in the blood and ∼80% hCD45^+^ in the bone marrow and spleen. Reported engraftment efficiencies for NSG mice preconditioned with busulfan at 15 weeks post CD34^+^ engraftment have been ∼10-40% in the blood and ∼60-80% in the bone marrow and spleen (Kang et al. 2016). We also found as reported by others a CD4:CD8 ratio of 3:1, and number of innate immune cells including NK cells and myeloid cells at a low percent of hCD45^+^ cells (1 to 0.5 percent NK cells, and < 5% myeloid lineage cells) in the blood and spleen. These results confirm that the NSG background does not support efficient innate immune reconstitution.

Taking these initial studies, a step further we show that using busulfan allows us to effectively reconstitute the hIL15Tg-NSG mouse background, but not he SGM3-NSG background. With both models we did not observe any evidence of GVHD as monitored by loss in body weight and accompanying pathologies. Use of busulfan reconstituted elevated levels of hCD45^+^ levels in the blood (∼20% vs ∼15%) and similar levels (∼70%) in the bone marrow and spleen between NSG and hIL15Tg-NSG mice. As expected, we observed high levels ∼15% or ∼10% of mature NK cells (CD56^+^, CD16^+^) in hIL15Tg-NSG mice compared to NSG mice in both the blood and the spleen. A study of the NK cell maturation marker perforin showed a similar trend, with more perforin expressing NK cells in the blood and the spleen of hIL15Tg-NSG mice.

In contrast to our findings in the hIL15Tg-NSG background, our findings show that the number of hCD45^+^ immune cells in the blood, bone marrow, and spleen of SGM3-NSG mice is generally lower than in NSG mice. We did observe that SGM3-NSG mice exhibited a greater percentage of human granulocytic MDSC in the blood, and an overall shift of monocytic MDSC to that of granulocytic in all measured tissues. We also observed elevated numbers of cDC1 and cDC2 dendritic cells, but not Mo-DC in the SGM3 mice compared to the NSG mice. However the CD14^+^ monocytes made up a significantly lower absolute proportion of engrafted leukocytes in SGM3 mice compared to NSG mice. In opposed to our findings in humanized IL15Tg-NSG mice utilizing busulfan, SMG3-NSG mice do not demonstrate as significant outcomes in myeloid cells, and do not in general agree with the literature for SGM3 mice reconstituted by TBI (Coughlan et al., 2016).

Furthermore, we want to figure out how many isolated CD34^+^ stem cells can be used in humanized mice. In general, 1 × 10^5^ CD34^+^ cells are used in typical reconstitutions with hUBC CD34^+^ cells, producing a 50–75 percent reconstitution when TBI is used for pre-conditioning, with 15–20 percent of these cells lacking CD38 and accounting for the majority of the reconstituting activity, as demonstrated by a similar reconstitution of 2 × 10^4^ to 1 × 10^4^ CD34^+^ cells (Drake et al., 2012). Ng Chee Ping stated that seven weeks after transplantation, the reconstitution limited their ability to identify human cells at the lowest cell dosage (300 cells). Compared with 10 weeks after transplantation, there are no significant variations in the total number of human cells found. Human hematopoietic cells were effectively found in the peripheral blood of mice receiving the most significant transplant cell dosages (10,000 progenitors) after 4 weeks (Ng Chee Ping, 2013).

The chimerism was found 3 weeks earlier than in animals implanted with lower progenitor cell dosages (Ng Chee Ping, 2013). Similarly, our result illustrates that at week eight post-transplantation, the average hCD45^+^ cells in hIL15Tg-NSG mice were high in mice engrafted with 1 × 10^5^ CD34^+^ stem cells compared with mice receiving 3 × 10^4^ and 1 × 10^4^ CD34^+^ stem cells. Interestingly, the engraftment % did not improve dramatically over the course of 8 weeks. This suggests for the hIL15Tg-NSG background that at least 1 × 10^5^ CD34^+^ cells are needed to ensure acceptable levels of engraftment.

In conclusions, this simple strategy for humanizing mice indicated that using the busulfan conditioning approach to humanize specialized NSG strains can result in the engraftment of human NK cells in the hIL15Tg-NSG background but is not an effective means to achieve high levels of engraftment in the SGM3-NSG background.

## ACKNOWEDGMENTS

The authors would like to acknowledge Rebecca Martin and Julie Farnsworth and the flow cytometry facility for help with processing and analysis of flow cytometry data. Funding for the research was obtained from the Virginia Commonwealth University School of Medicine, Virginia Commonwealth University Office of Research, and the Massey Cancer Center.

---

**Figure S1.**
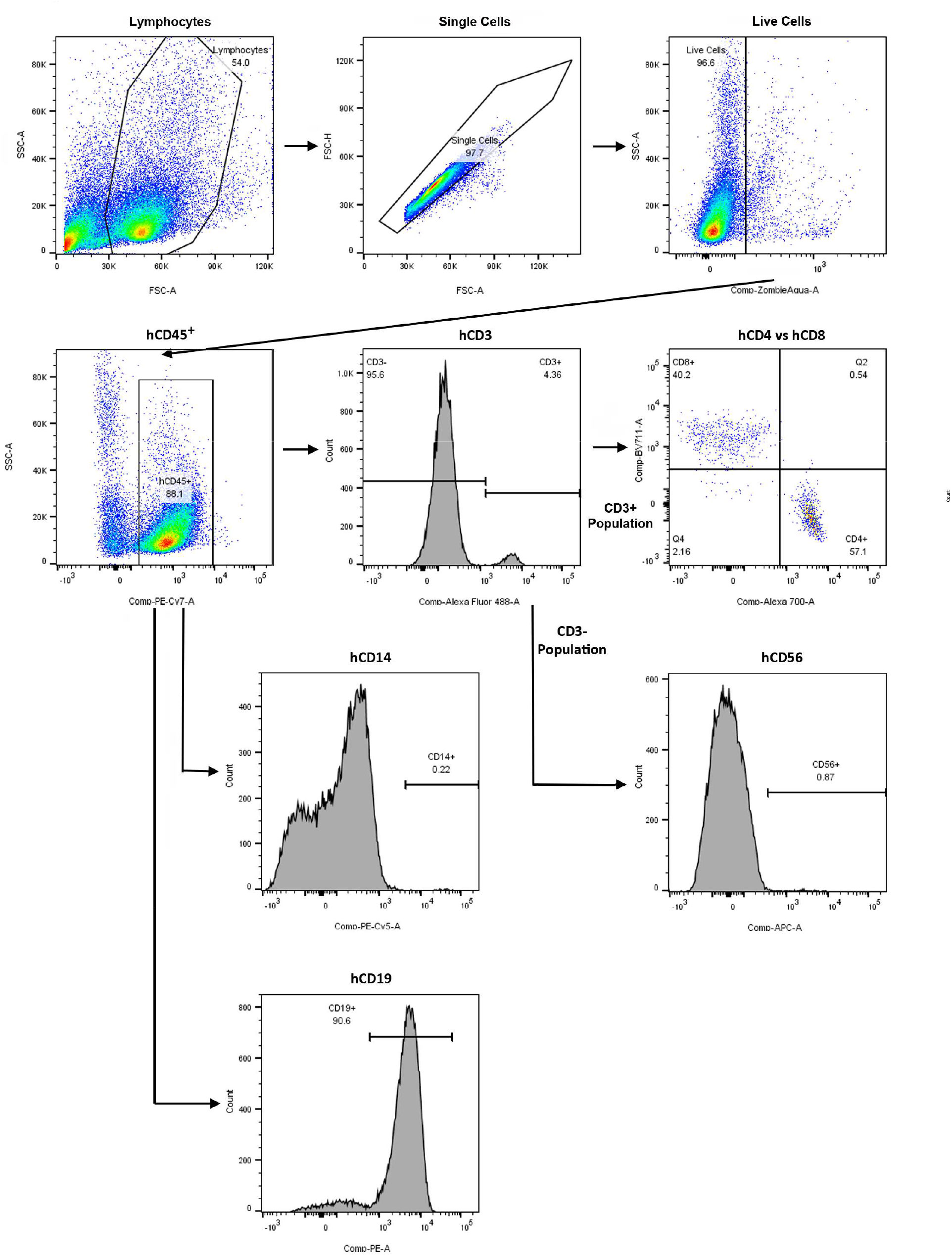
Gating Strategy for the Identification of Human Immune Cell Populations in Humanized NSG Mice. Representative gating strategy for the identification of live hCD45^+^ leukocytes, CD3^+^ CD4^+^ or CD3^+^ CD8^+^ human T cells, CD3^-^ CD56^+^ human NK cells, hCD14^+^ human monocytes and hCD19^+^ human B cells.

**Figure S2.**
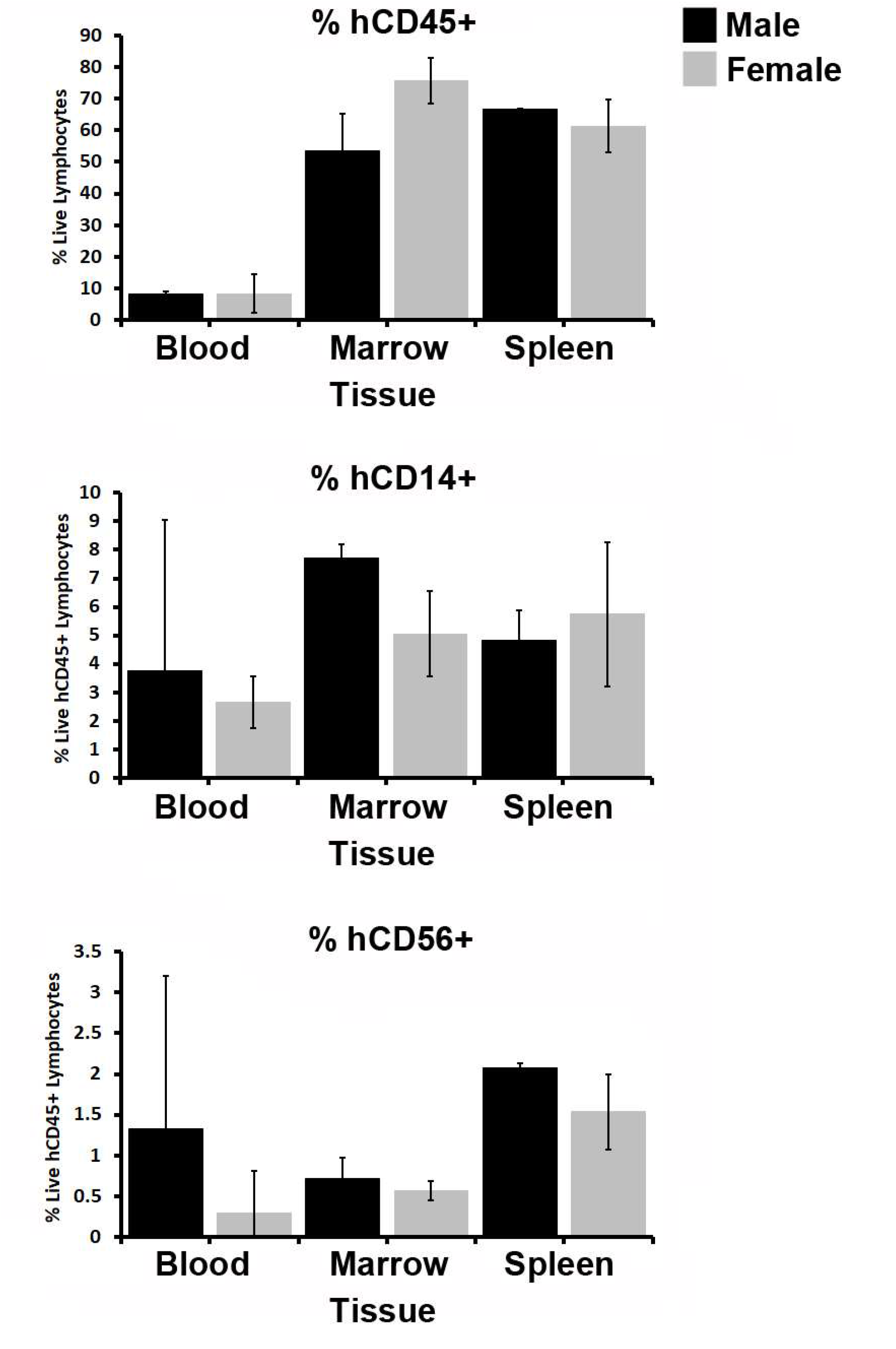
Analysis of Sex as a Variable in Humanization of NSG mice. The blood, spleen and bone marrow of female or male NSG mice 15 weeks post transplantation with congruent female or male hUCB-derived CD34^+^ cells used were analyzed for abundance of hCD45^+^ human leukocytes, CD56^+^ human NK cells, and hCD14^+^ human monocytes.

**Figure S3.**
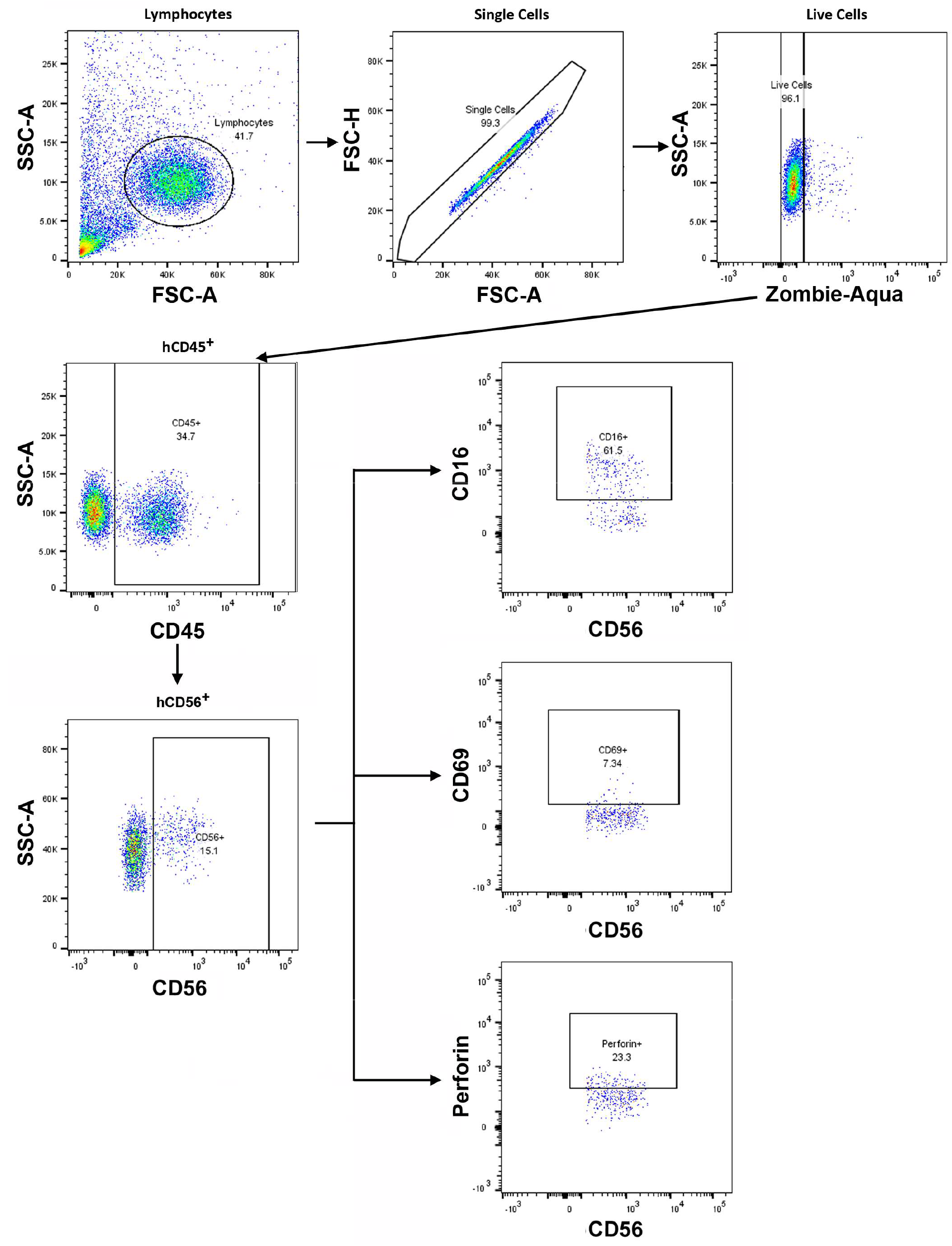
Gating Strategy for the Identification of Human Immune Cell Populations in Humanized hIL15-Tg-NSG Mice. Representative gating strategy for the identification of live hCD45^+^ leukocytes, CD56^+^ CD16^+^ or CD56^+^ CD69^+^ or CD56^+^ perforin^+^ human NK cells.

**Figure S4.**
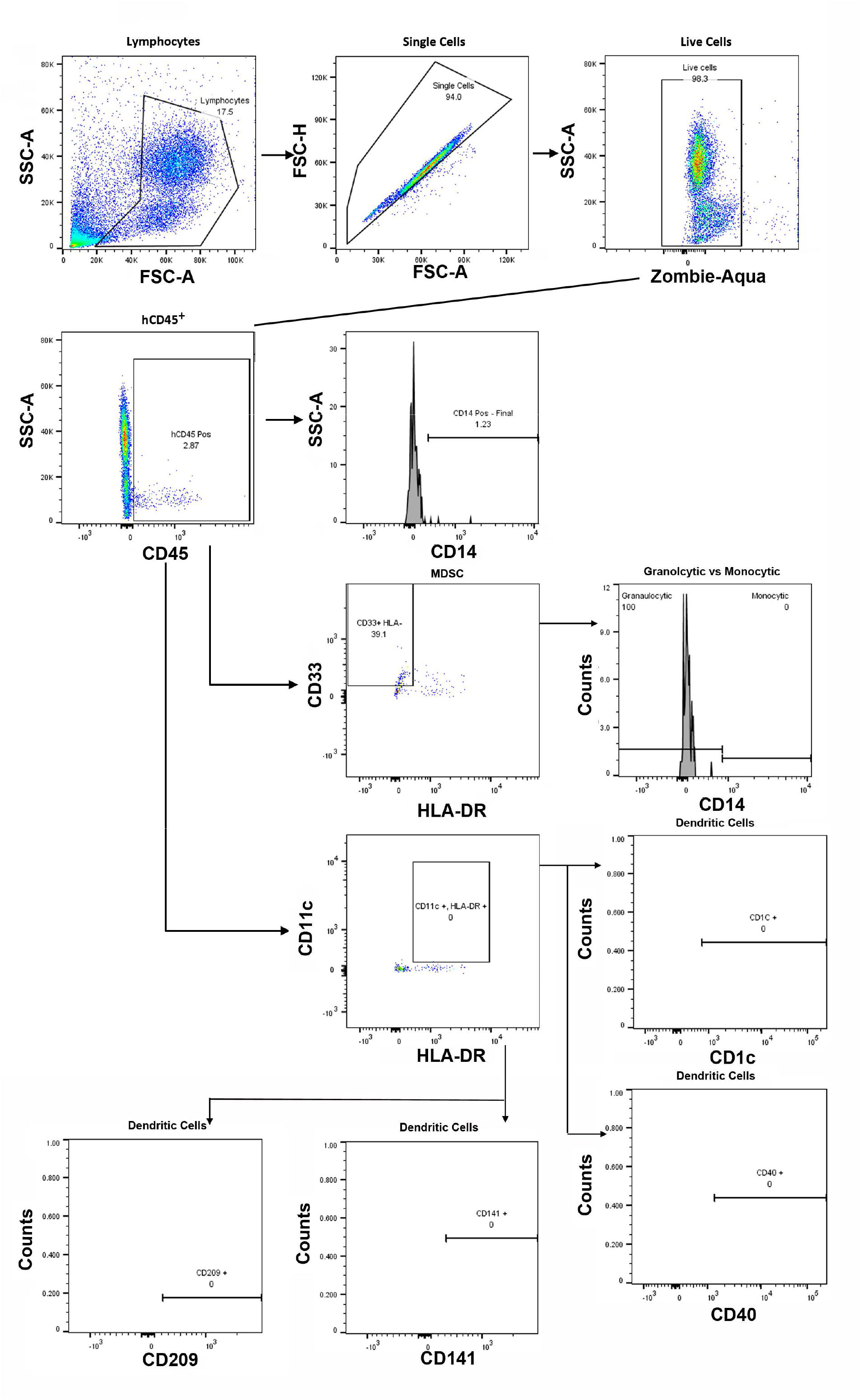
Gating Strategy for the Identification of Human Immune Cell Populations in Humanized SGM3-NSG Mice. Representative gating strategy for the identification of live hCD45^+^ leukocytes, CD33^+^ HLA^-^ MDSC and the CD14^+^ granulocytic and CD14^-^ monocytic populations, CD11c^+^ HLA-DR^+^ dendritic cells and the CD1c, CD141, CD209 dendritic cell populations and the CD40 activation status.

